# Optimising biomedical relationship extraction with BioBERT

**DOI:** 10.1101/2020.09.01.277277

**Authors:** Oliver Giles, Anneli Karlsson, Spyroula Masiala, Simon White, Gianni Cesareni, Livia Perfetto, Joe Mullen, Michael Hughes, Lee Harland, James Malone

**Affiliations:** SciBite, Wellcome Genome Campus, Hinxton, United Kingdom; Department of Biology, University of Rome Tor Vergata, Rome, Italy; European Bioinformatics Institute, European Molecular Biology Laboratory, Wellcome Genome Campus, Hinxton, United Kingdom

## Abstract

Text mining is widely used within the life sciences as an evidence stream for inferring relationships between biological entities. In most cases, conventional string matching is used to identify cooccurrences of given entities within sentences. This limits the utility of text mining results, as they tend to contain significant noise due to weak inclusion criteria. We show that, in the indicative case of protein-protein interactions (PPIs), the majority of sentences containing cooccurrences (∽75%) do not describe any causal relationship. We further demonstrate the feasibility of fine tuning a strong domain-specific language model, BioBERT, to analyse sentences containing cooccurrences and accurately (F1 score: 88.95%) identify functional links between proteins. These strong results come in spite of the deep complexity of the language involved, which limits the accuracy even of expert curators. We establish guidelines for best practices in data creation to this end, including an examination of inter-annotator agreement, of semisupervision, and of rules based alternatives to manual curation, and explore the potential for downstream use of the model to accelerate curation of interactions in the SIGNOR database of causal protein interactions and the IntAct database of experimental evidence for physical protein interactions.

## Introduction

Recent years have seen the flourishing of deep learning based natural language processing. Transformer architectures [1] have enabled models to better understand the grammar and discourse of texts to the extent that they have become capable of inventing convincing blogs, social media posts and even more longform content like Wikipedia articles [2]. Alongside these architectural developments there has been a growing recognition of the value of transfer learning. With transfer learning, models are initially trained to perform automatable tasks on an enormous corpus of text. This task may involve, for example, predicting words that have been omitted from sections of text [3]. For such a task there is no need for curation since the training data can be automatically generated, allowing for exponential upscaling. These models can then be slightly modified, either in terms of their architecture or, more recently, simply in terms of instruction [4], to perform new tasks. When undertaking these specific tasks, the vast majority of the model remains as it was, and the general linguistic knowledge acquired through the initial training, including grammatical principles that it would be near impossible to learn from the relatively miniscule datasets used in the fine tuning stage, can be leveraged. As a result, it is now possible to achieve strong results with relatively little curation, driving down the previously prohibitive cost of developing state of the art models within specific domains. Furthermore, models with domain specific pretraining have also been developed, such as BioBERT [5], whereby the initial training is followed by further training on life science corpora such as MEDLINE and PubMedCentral, resulting in further gains on life science specific tasks. BioBERT was released with three fine-tuned variants of the base model for performing named entity recognition, question answering and relationship extraction. Research has shown a moderate improvement in the extraction of biomedical relationships using BioBERT as opposed to base BERT [6].

The sheer volume of publications in the life sciences presents a significant challenge to researchers attempting to monitor developments and compile the consolidated resources upon which modern research is largely dependent. The potential to streamline these operations by fine tuning state of the art deep learning models to extract causal relationships from text therefore warrants research [7].

Protein-protein interactions (PPIs) are critical to a great number of disciplines including the established, like pharmacy, and the up and coming, like synthetic biology [8]. They are often described using deeply complex and technical language and, as the majority of sentences containing multiple proteins do not describe any interaction, they are resistant to extraction with simple, traditional text mining methods. The extraction of PPIs from text is therefore a pivotal problem to be solved, and empowering researchers to more effectively collect these data may have significant real world benefits in downstream applications. We therefore chose to focus on this particular task, but the methods described are largely transferable to the extraction of other relationships from text. As the state of the art in natural language processing has progressed over the past years, these methods have been applied to PPI extraction from text [9–11]. However, different use cases define interactions differently, which can be seen by the number of papers [11–13] which achieve disparate results on the major datasets of BioInfer [14] and AIMed [15]. For example, either causal or physical interactions may be sought, or both. There may also be other criteria for extracted sentences, such as requiring that they describe novel discoveries. As models continue to become more capable of abstracting patterns from fewer samples of data, the feasibility of creating highly specific models for niche concerns increases, and the onus of generating data shifts away from consortia and towards individuals and organisations. It is therefore key to have clear guidelines for best practices in this data creation.

In this paper, we demonstrate the feasibility of using traditional text mining as a starting point to narrow down a curation space to facilitate more rapid generation of datasets, and also to narrow down the execution space once the model is trained. We then assess whether there is a need for inter-annotator agreement in the curation of data, and whether semisupervision can reduce curation time, and suggest best practices based on the outcomes. We proceed to compare the efficacy of rules-based methods, deep learning methods bootstrapped with data collected via rules-based methods, and deep learning methods trained on curated data. Finally, we outline some initial testing undertaken to assess the value added to current PPI database curation methods both with and without further task-specific fine tuning.

## Materials and methods

### Precuration

Curation is a time consuming and thereby costly process. Without appropriate data preparation, an individual curator can read only a handful of papers per hour, which may or may not contain examples of interest. In order to narrow down the curation space for our tasks, we defined high level minimum criteria. It is rare that a text describes an interaction between two different proteins without containing the explicit mention of both proteins. We used named entity recognition (NER) software, TERMite, to tag life science entities within papers drawn from MEDLINE. This enabled us to extract only sentences containing relevant, high level patterns. In the case of PPIs, we looked for two or more gene hits, aligned to HGNC [16], within a given sentence. In auxiliary tests for semisupervision we also looked at sentences containing pairs of drug and adverse event hits, aligned to ChEMBL [17] and MeDDRA [18] respectively, and pairs of drug and gene hits. Filtering sentences in this way not only allows for more rapid curation but also results in fewer sentences being required for training, as the model only needs to comprehend the nuances of a niche subset of sentences. This same step can be used at inference such that only sentences of interest are passed to the model, resulting in significant savings of compute and time.

### Data preparation

#### Manual curation and Inter-annotator agreement

To assess the impact of inter-annotator agreement, three curators independently curated an initial set of 925 sentences, taken from MEDLINE, within which our NER system had identified two or more genes/proteins. Experienced curators attempted to identify PPIs according to criteria provided (see Table 1, or S1 Appendix for full detail). Our criteria were aimed at developing a recall-oriented model which could potentially then be further fine tuned to more specific use-case criteria as required. Concordance between all three curators was observed in 451 of 925 sentences (48.8%), and concordance between at least two of three curators was observed in 889 of 925 sentences (96.1%). A subset of 170 sentences enjoyed agreement from two curators while the third curator had indicated they were unsure as to the correct category. We curated these sentences with a fourth curator and found agreement with the 2:1 majority in 155 cases (88.2%).

**Table 1.** Classes of protein-protein sentences.

| Label | Class | Example |
| --- | --- | --- |
| C | Coincidental mention | A and B were measured. |
| P | Positive | A binds to B. |
| N | Negative | A does not bind to B. |
| I | Incorrect entity recognition | Turn to PAGE1 to read about A. |
| ? | Don't know/unclear |  |

Each sentence was labelled with a single high level classification from the table above. Further clarification about edge cases was provided when necessary, with an overarching goal to identify the broadest sense of PPIs.

The high level of disagreement amongst curators (with at least one curator dissenting in 51.2%of cases), illustrates the complexity of the problem even as approached by human experts. We also observed that the number of sentences deemed to contain coincidental mentions of genes significantly outnumbered the number of sentences deemed to describe interactions, illustrating the need for models to differentiate between these classes in order to reliably automate the identification of PPIs within literature.

Although two curators could process more sentences than three in a given number of person-hours, the resulting number of sentences with agreement between two annotators per unit time was similar with either three curators or two curators. In our case, we found inter-annotator agreement between at least two of three curators in 96.1%of sentences, as opposed to a mean average of 64%between the possible pairings of two curators. If three curators curate n sentences in t person-hours, we would expect two curators to curate 1.5n sentences in t, and 0.64 · 1.5n ≈ 0.961n (or 0.96 ≈ 0.961). It should be noted that agreement between two out of two curators represents a concordance rate of 1, whereas agreement between two out of three curators represents concordance of 0.67. We determined to proceed with three annotators to allow us to assess the efficacy of models trained with varying degrees of concordance and to get the strongest possible gold standard set.

#### Semisupervision

The class imbalance between coincidental and positive mentions lowers the cost efficiency of curation. To improve the prevalence of positive sentences we utilised the background knowledge from StringDB [19], a public database of mined PPIs, and repeated the curation step after filtering the sentences such that only those which contained two genes identified to have a strong likelihood of interacting, signified by a StringDB combined score of *≥* 950, were retained.

Despite this additional data preparation step, we observed negligible differences in the rate of identification of positive interactions. Moreover, after filtering for gene pairs with high combined scores in StringDB, all three annotators agreed on interactions within only 118 of 1085 sentences (10.9%) and two of three annotators agreed on interactions in a further 136 sentences, bringing the combined total to 254 (23.4%). In the randomly selected set of sentences, these figures were 15.2%and 26.1%respectively. More broadly, we observed similar rates of agreement to the initial set of sentences with all three annotators agreeing in 51.3%of cases (compared to 48.8%in the initial set) and two out of three annotators agreeing in 94.9%of cases (compared to 96.1%). The one clearly observable difference was a marked increase in the identification of sentences containing coincidental mentions (increasing from 47.8%/25.3%to 58.7%/35.2%with agreement between two and three curators respectively).

We repeated this with even stricter StringDB combined scores of *≥* 995, and once again found no improvement in the rate of identification of positive interactions. This may be a result of well established interactions being assumed knowledge and therefore rarely being explicitly stated. We therefore continued using the initial randomly selected set of sentences for data preparation.

To assess whether semisupervision might be more applicable in the case of a different relationship, we attempted to apply a similar methodology to drug/adverse reaction pairs. Adverse reactions are difficult to identify using traditional string matching methods as they are lexically identical to non-adverse reaction indications. As such, a sentence which appears to contain a drug and an adverse reaction may in fact describe the opposite, with the drug being described as a treatment for the indication in question. This indicates there is potential for the use of semisupervision in the development of models to extract this relationship.

We used TERMite to identify indications mentioned on FDA drug labels within the warnings section, and considered these indications to be adverse reactions caused by the drug to which the label belonged. We proceeded to curate 100 sentences containing this subset of drug/indication pairs and a further 100 sentences containing randomly selected drug/indication pairs, as identified by our NER system in MEDLINE. In both the semisupervised and the randomly selected set, 23 sentences were deemed to likely describe a drug causing an adverse event. It is very common for a drug to list the indication it treats as a side effect (e.g. headaches being a side effect of aspirin), so we postulated that one possible way to improve on this result would be to exclude any indication mentioned on the drug label which is also known to be a condition treatable with said drug. Repeating the above methodology but excluding approved treatments, as listed in ChEMBL, resulted in a minor improvement of 28 likely positive sentences being identified in the 100 curated.

We undertook one final round using ChEMBL to identify drug-gene pairs wherein the gene was a known target of the drug. In this case we found that 58.1%of randomly selected sentences containing a drug and a gene likely described a targeting relationship. In the semisupervised set, this rose to 89.9%. In conclusion, semisupervision may provide a valuable means to increase the ratio of sentences containing the desired relationship to those containing coincidental mentions, but this value is case dependent.

#### Final curation

We continued with curation using randomly selected sentences containing two genes/proteins, with no further filtration, ultimately collecting 1408 sentences deemed by at least two curators to contain an interaction, of which 308 were randomly selected and allocated to a test set. These sentences were combined with a correspondent number of sentences deemed by at least two curators to contain coincidental mentions to create our primary training and testing sets.

### Approaches

#### Rules-based approaches

We attempted to use some simple rules-based approaches to provide baselines for deep learning comparisons. The most reliable rule we could identify was to look for two genes/proteins alongside a bioverb indicative of an interaction. We compiled a list of 41 such bioverbs (S2 Appendix). In an attempt to improve precision, we also used a second method utilising the natural language processing library spaCy [20] to generate dependency trees of our sentences in order to confirm that the genes identified were grammatical children of the bioverb identified. If the bioverb did not appear to link the two genes, these sentences were considered to be coincidental as opposed to positive. Curators did observe a number of other potential rules such as excluding genes separated by commas in lists, and excluding common pairs such as BRCA1/BRCA2 and CD3+/CD4+. Each of these rules excludes both positive and coincidental examples (albeit not in equal numbers). For the purposes of this experiment we decided to keep the number of rules to a minimum to avoid any confounding effects. Nevertheless, a more sophisticated ruleset may improve results if care is taken to balance precision and recall.

#### Mixed approach

To define deep learning baselines, we first attempted to train a model using data collected with the rules-based methods defined above. The risk inherent in using rules-based approaches for data gathering is the introduction of bias, as one is effectively training a model to recognise one’s simplified ruleset, as opposed to exposing it to a truly representative sample of the more nuanced relationship you hope to identify. To assess whether BioBERT was abstracting patterns from our rules-generated training sets, we removed sentences containing a subset of five bioverbs from our training set and used these as our test set. Training was then performed on sentences containing the remaining set of bioverbs, after replacing any gene hits identified by TERMite with a normalised token. We observed F1 accuracy of 0.584 on unseen bioverbs using the basic method, and F1 accuracy of 0.801 on unseen bioverbs using the spaCy method. This indicates that, while both methods led to some degree of abstraction, this was much more pronounced in the method using grammatical dependency parsing than the method using simple term identification. This insight may be useful if curated data is not available.

#### Deep learning approach

Using the same parameters as in the mixed approach, we trained BioBERT on our manually curated sentences.

### In-the-wild testing

Following encouraging results from a preliminary examination of the model’s output, curators at SIGNOR [21] and IntAct [22] examined two sets of data extracted from the CORD19 dataset [23]. A small, custom vocabulary of PPI measurement techniques was used to filter the documents (S3 Appendix), and pairs already represented in the SIGNOR database were excluded. One set was created by identifying sentences with two proteins using TERMite. The second set used the model prediction as an additional filter. Another round of curation was then undertaken to assess coverage of particular genes of interest in which TERMite was used to identify proteins listed by SIGNOR/IntAct as being high priority.

## Results

From our results in Table 2, it is clear that the direct application of our simple, rules-based methods had limited effect in further filtering sentences identified as containing two proteins. However, despite a relatively low F1 accuracy of 0.465 from the most basic ruleset, which simply looked for sentences with two genes/proteins and one molecular bioverb, the precision of 0.666 provides a sufficient weighting of positive sentences to allow BioBERT to abstract common patterns, resulting in significant improvements when a model is trained on sentences identified by the ruleset (F1 accuracy 0.761, an increase of almost +0.3). Offsetting this, the relatively high number of incorrect sentences identified by the ruleset limits the final accuracy of the model. We attempted to address this by using dependency parsing to increase our confidence that the two proteins mentioned in a sentence were in fact linked by the molecular bioverb we had identified. When directly applied, this second ruleset did result in notably higher precision (0.759, compared to 0.666). Unfortunately, this came at a significant cost to recall which fell from 0.357 in the original ruleset to 0.133, ultimately resulting in a much reduced F1 accuracy of 0.227. Likewise, training BioBERT on sentences identified by this second ruleset resulted in higher precision (+4.6%) and lower recall (−10.4%) than the model trained on the original ruleset, and an overall lower F1 accuracy of 0.734 (−2.7%). One point of interest is the dramatic rise in recall from the pure rules classification (0.133) to the BioBERT model trained on the output of those rules (0.708). This may indicate the strength of BioBERT in conceptual abstraction, where a simpler model would simply look for similar keywords, resulting in recall more similar to that of the original ruleset. It also correlates with our prior finding that using dependency parsing to generate a training set results in better performance on sentences with unseen key words.

**Table 2.**
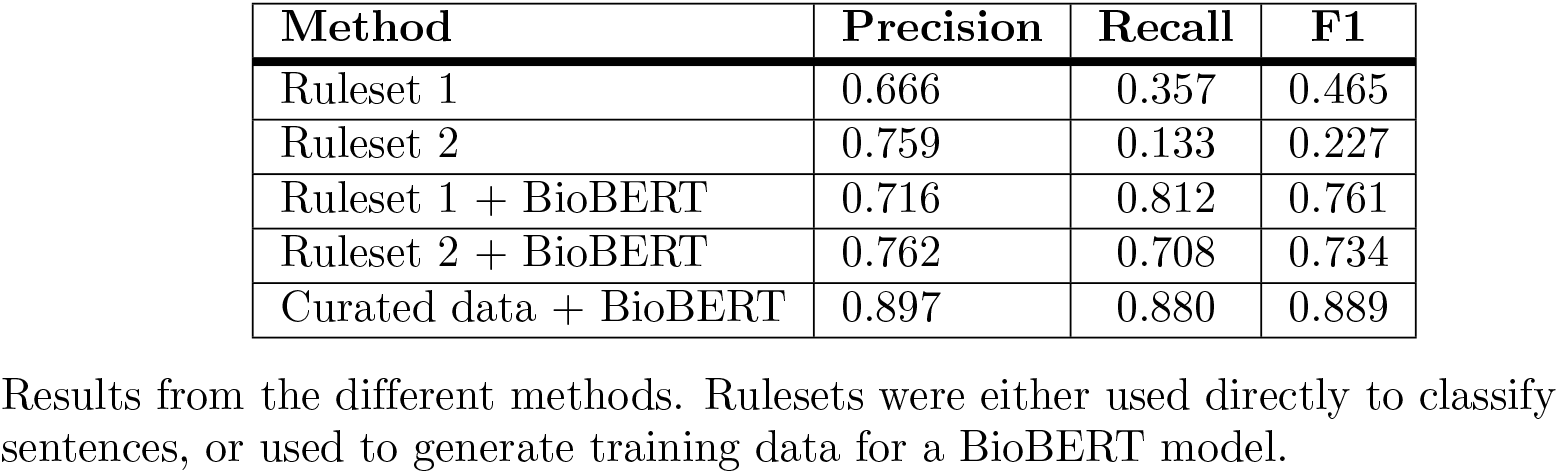
Method results.

Broad rules tend to result in a trade off between precision and recall, while highly specific rules are challenging to identify and can easily result in overfitting. The promise of deep learning lies in its capability to recognise nuance and patterns, which are often not easily abstracted into explicit rules, through exposure to real world data. Relative to both the rules-based approaches and the deep learning approaches bootstrapped on data from those rules-based approaches, the deep learning model trained on curated data was significantly stronger. Furthermore, after training BioBERT with fractional subsets of our final curated data, we observed that these results could be achieved with relatively few examples, with F1 accuracy exceeding 0.8 having only seen 100 examples per class and exceeding 0.85 having seen just over 300.

In order to get an indication as to whether different tasks would require similar amounts of training data, we repeated this with the Genetic Association Database (GAD) [24] dataset for gene/disease associations. We observed similar accuracy curves when plotting results from different tasks (Fig 1). 95%of the peak accuracy was achieved after 250 sentences for the PPI model, and after 400 sentences in the gene-disease association model. Rounding up, 500 sentences per class may be a useful benchmark for cost efficiency for incrementally training BioBERT or similar language models. The different tasks do achieve different levels of accuracy, which may be the result of different levels of stringency during curation or different complexities in the language describing the different relationships.

**Fig 1.**
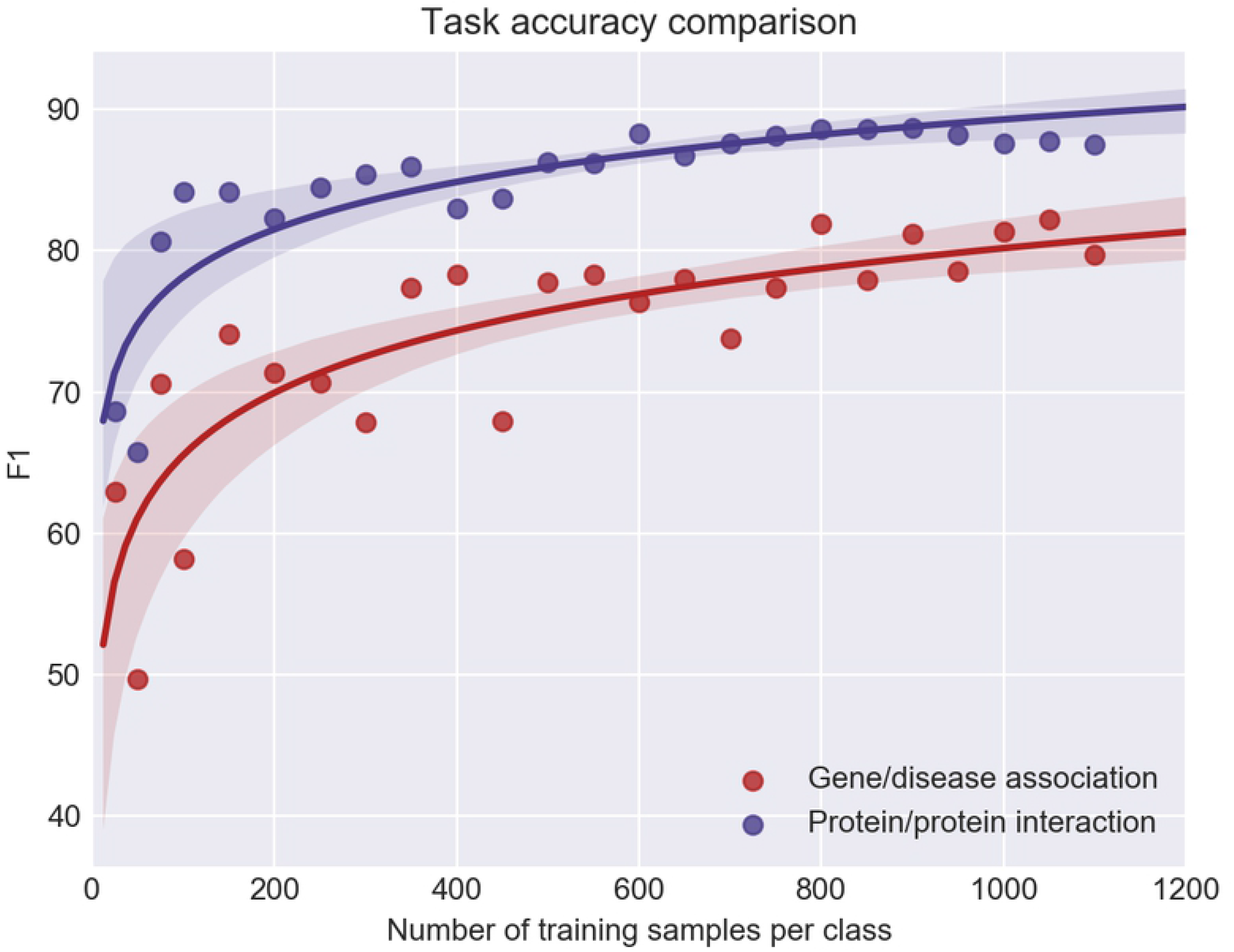
Task accuracy curves. Comparison of the relationship between model F1 accuracy and number of training samples per class in different relationship extraction tasks.

**Fig 2.**
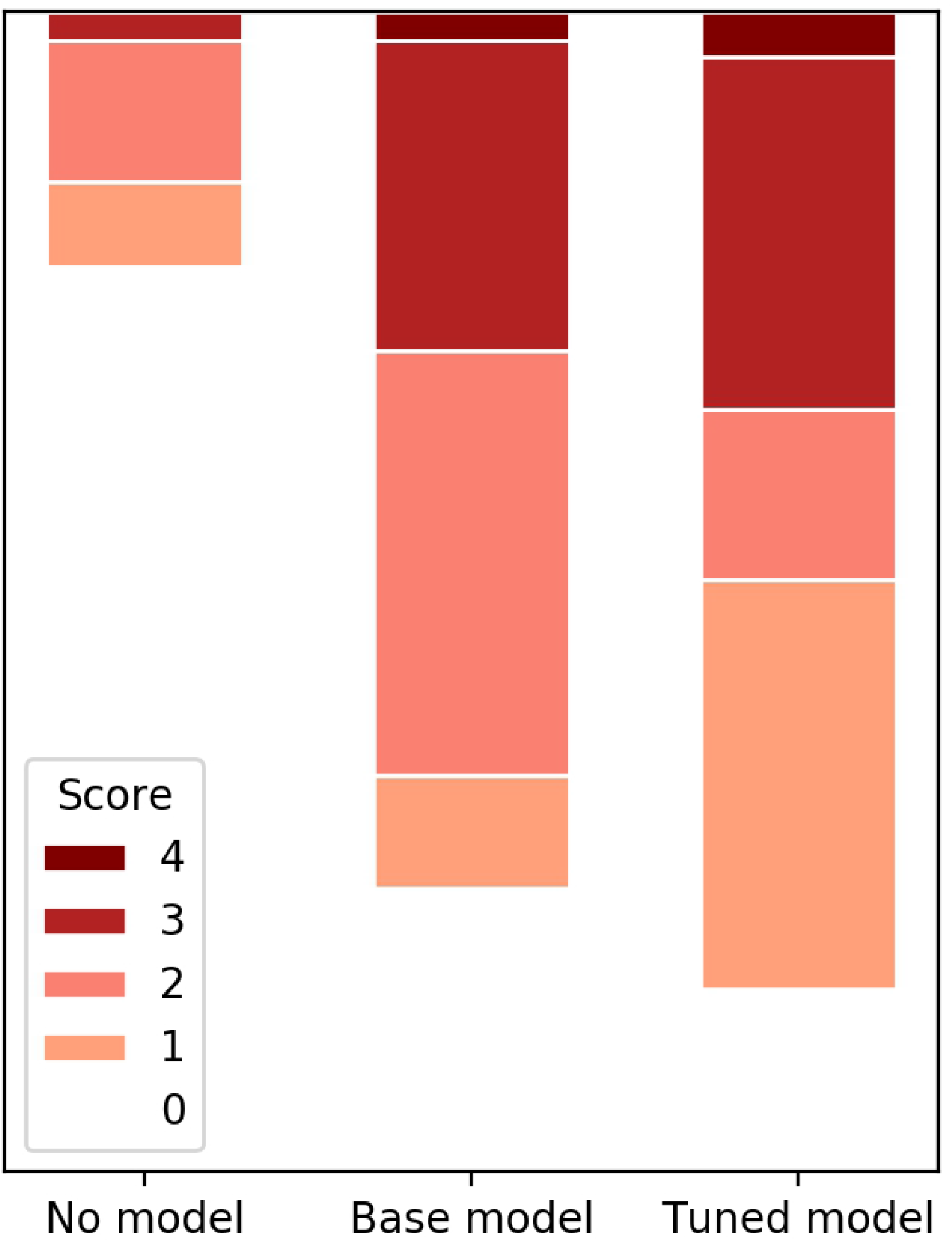
SIGNOR/IntAct sentence scoring. Comparison of the distribution of curation scores with either no model as a filter, the original model as a filter, or the model tuned to the curation criteria of SIGNOR/IntAct as a filter. In each case, sentences contained two genes/proteins and were extracted from documents taken from the CORD19 dataset which contained one or more mentions of a PPI measurement technology.

During curation, it can be challenging to ascertain whether an interaction is being described from reading a single sentence. Rather than forcing difficult sentences into an existing category, we decided to collect these sentences separately. We hypothesised that having a bin for sentences which did not clearly belong in either the coincidental bin or the positive bin may enable us to train models with an emphasis on either precision or recall, with each model suited to different use cases. To verify this, we replaced 20%of positive sentences with sentences labelled as unclear/unknown by at least two curators. Training a model on this new dataset and testing on the original test set resulted in recall of 0.883 (+0.003) and precision of 0.866 (−0.031), with an F1 score of 0.875 (−0.014). Replacing 20%of coincidental sentences resulted in recall of 0.857 (−0.023) and precision of 0.895 (−0.0025), with an F1 score of 0.876 (−0.013). Although the general trend was as expected, the introduction of these difficult sentences resulted in slight accuracy reductions across the board as compared to baseline, excepting a miniscule increase in recall when the unclear/unknown sentences were introduced into the positive set.

Contributors to SIGNOR, a database of causal relationships between biological entities, and to IntAct, a database that captures experimental evidence supporting physical interactions, examined two sets of sentences. Both sets of sentences were taken from documents containing mentions of PPI measurement technologies within the CORD19 dataset. Named entity recognition software was used to identify sentences containing two genes/proteins. These sentences were used, as is, to create a control set, and a second set was created by feeding sentences to the trained model and only retaining those predicted to contain an interaction. The sentences were curated by individuals from SIGNOR and IntAct according to the criteria in S4 Appendix. These criteria were applied to both causal interactions and physical interactions. We took the highest of these two scores to be the score of the sentence. 31/40 (77.5%) of sentences in the set within which the model had identified a PPI were deemed to describe an interaction at some level, as opposed to 9/40 (22.5%) in the sentences without any model filtering. Although 77.5%falls short of the model’s accuracy on the curated test set, it should be noted that the test set was created using the majority opinion of three independent curators, and the 77.5%concordance between the model predictions and the SIGNOR/IntAct curators exceeds the average 64%concordance observed between pairs of curators. Some illustrative errors are collated in Table 3. The criteria for SIGNOR/IntAct also differ from the initial curation criteria, erring towards being rather stricter. Illustrative examples of these differences can be found in Table 4 The extraction of sentences was deemed more efficient than searching and reading entire papers, and using the model as an additional filter resulted in a much higher rate of useful sentences.

**Table 3.** Illustrative incorrect model predictions.

| Example | Model prediction | SIGNOR/IntAct score | Comment |
| --- | --- | --- | --- |
| We also demonstrate that the antiaging effect of Sip2 acetylation is independent of nutrient availability and TORC1 activity. | Positive | 0 | Sentences denying an interaction were extremely rare in the initial training data. All three curators agreed on only one negative example. |
| This comprises TRF1 and TRF2 which directly bind the duplex structure and POT1 which interacts with the single-stranded overhang tail. | Positive | 0 | Binding and interacting, but with a non-protein molecule. |
| We constructed and delivered the shRNA-resistant myc-tagged DDX1 expression plasmids into the DBT cells, within which the endogenous DDX1 had already been knocked down by using shDdx1-1 ( Figure 6A, lanes 4-6) . | Positive | 0 | An edge case - DDX1 is tagged with myc, but via recombinant DNA technology as opposed to any interaction between independent proteins. |
| Christopher Stroh (Muenster, Germany) presented an intriguing strategy for enhancing apoptosis sensitivity of tumor cells by transfection of the NF-kB inhibitor IκBa fused with the viral protein VP22... | Positive | 0 | Another edge case - fusion proteins do not interact as independent entities. |
| MICAL1 colocalizes with Rab8a. | Positive | 0 | Colocalisations are not considered positive according to either curation protocol. |

Many of the models mistakes are understandable and may be addressed with targeted improvements to the training data.

**Table 4.**
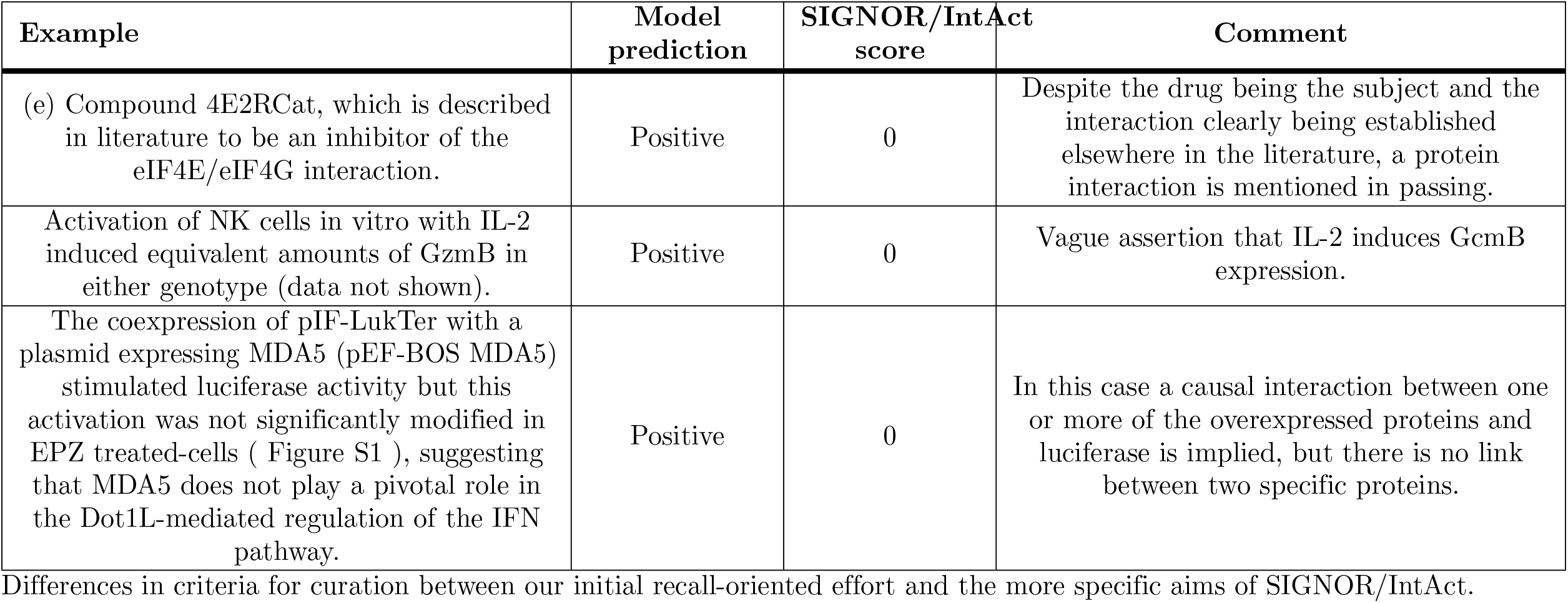
Differences in curation criteria.

| Example | Model prediction | SIGNOR/IntAct score | Comment |
| --- | --- | --- | --- |
| (e) Compound 4E2RCat, which is described in literature to be an inhibitor of the eIF4E/eIF4G interaction. | Positive | 0 | Despite the drug being the subject and the interaction clearly being established elsewhere in the literature, a protein interaction is mentioned in passing. |
| Activation of NK cells in vitro with IL-2 induced equivalent amounts of GzmB in either genotype (data not shown). | Positive | 0 | Vague assertion that IL-2 induces GzmB expression. |
| The coexpression of pIF-LukTer with a plasmid expressing MDA5 (pEF-BOS MDA5) stimulated luciferase activity but this activation was not significantly modified in EPZ treated-cells ( Figure S1 ), suggesting that MDA5 does not play a pivotal role in the Dot1L-mediated regulation of the IFN pathway. | Positive | 0 | In this case a causal interaction between one or more of the overexpressed proteins and luciferase is implied, but there is no link between two specific proteins. |

Differences in criteria for curation between our initial recall-oriented effort and the more specific aims of SIGNOR/IntAct.

It was deemed that the potential existed for the model to improve curation time at SIGNOR/IntAct, so a further test was carried out using a specific set of proteins of interest. TERMite was used to identify sentences containing two proteins, at least one of which was listed by SIGNOR/IntAct as being of interest. These sentences were ordered according to the proteins present such that all sentences supporting an interaction between a pair of proteins could be considered collectively. 144 of 210 (68.6%) sentences were deemed to contain an interaction. The accuracy was likely impacted by the ordering, with certain pairs posing particular difficulty for the model. One example of this was the pair IL17 and IL17R, often captured together as ‘IL17/IL17R’ after the same fashion as complexes that the model has been trained to consider positive (for example, ‘SWI/SNF complex’ or ‘Mre11/Rad50 complex’). Certain other trends were apparent in the negative sentences, such as the capturing of colocalisations, and of interactions between proteins and non-protein molecules, such as RNAs. Further work might involve targeted curation to address these observable trends. Gains may also be made in other ways. For example, databases like SIGNOR and IntAct aim to capture evidence of an interaction. In this case, sentences which simply assert an interaction takes place based on prior work have less value. A second model to classify the novelty of an assertion might help to further filter sentences to meet their criteria.

We endeavoured to explore the efficacy of further fine tuning our trained model, using the higher scoring sentences from the above curation as positives and the lower scoring sentences as negatives, in an attempt to increase the number of sentences output by the model that would be immediately curatable for inclusion in SIGNOR/IntAct. The model developed above was fine tuned on a further 47 sentences per class where sentences scored 3/4 in the previous rounds of curation were considered positive, and sentences scored 0/1/2 were considered negative. Sentences were then extracted from the CORD19 dataset using the protocol described above. 84.31%of 102 sentences curated were deemed to contain some level of interaction according to the criteria of SIGNOR/IntAct, an increase of 8.7%. A significant number of the remaining mistakes were interpretable, such as interactions between one protein and one gene (see Table 5). The number of sentences scoring either 3 or 4 increased from 29.27%to 34.31%(+5.04%). Examples can be seen in Table 6. These results were positive, especially considering the low number of training samples available. One caveat to note is that this more stringent model was notably less likely to make positive predictions, so recall was likely reduced. The importance of recall depends largely on the ambition of the curators. For example, a database with low coverage and an aim to increase this indiscriminately and quickly will benefit from a model with a focus on precision. Inversely, a database with significant coverage or with an aim to target specific entities will benefit from recall to ensure those few positives not already represented within the database are not missed. This once again illustrates the importance of models being amenable to fine tuning, as even within one database models with different emphases may be required at different stages of its life cycle.

**Table 5.** Illustrative incorrect model predictions.

| Example | Model prediction | SIGNOR/IntAct score | Comment |
| --- | --- | --- | --- |
| In addition, no HDAC6-derived phosphopeptide was detected in our analysis suggesting that GRK2 does not exert its proviral role through HDAC6. | Positive | 0 | Negative examples still pose a problem for the model, as they remain poorly represented in the training data. |
| ChIP experiments confirmed that there was a strong association of STAT3 with the GFAP promoter, suggesting the existence of mechanism that facilitates access of the STAT3 complex to the GFAP promoter. | Positive | 0 | Interaction between protein and gene. |
| To evaluate whether TRIM25 could counteract the reduction of the antiviral response mediated by Dot1L inhibition, TRIM25 overexpression experiments were carried out. | Positive | 0 | Hypothesis positing a causal interaction. |
| Depletion of STT3A, but not STT3B, causes a modest induction of the unfolded protein response (UPR) pathway. | Positive | 0 | Pathway induction implies an interaction may exist but none is explicitly described. |
| (50) found that WNT5a can increase fibroblast proliferation through a "noncanonical" or b-catenin/TCF-independent signaling mechanism, indicating that both canonical and noncanonical WNTs may contribute to tumorigenesis. | Positive | 0 | Effectively another negative example, asserts an unspecified signalling mechanism independent of protein mentioned. |

Many of the models mistakes are understandable and may be addressed with targeted improvements to the training data.

**Table 6.**
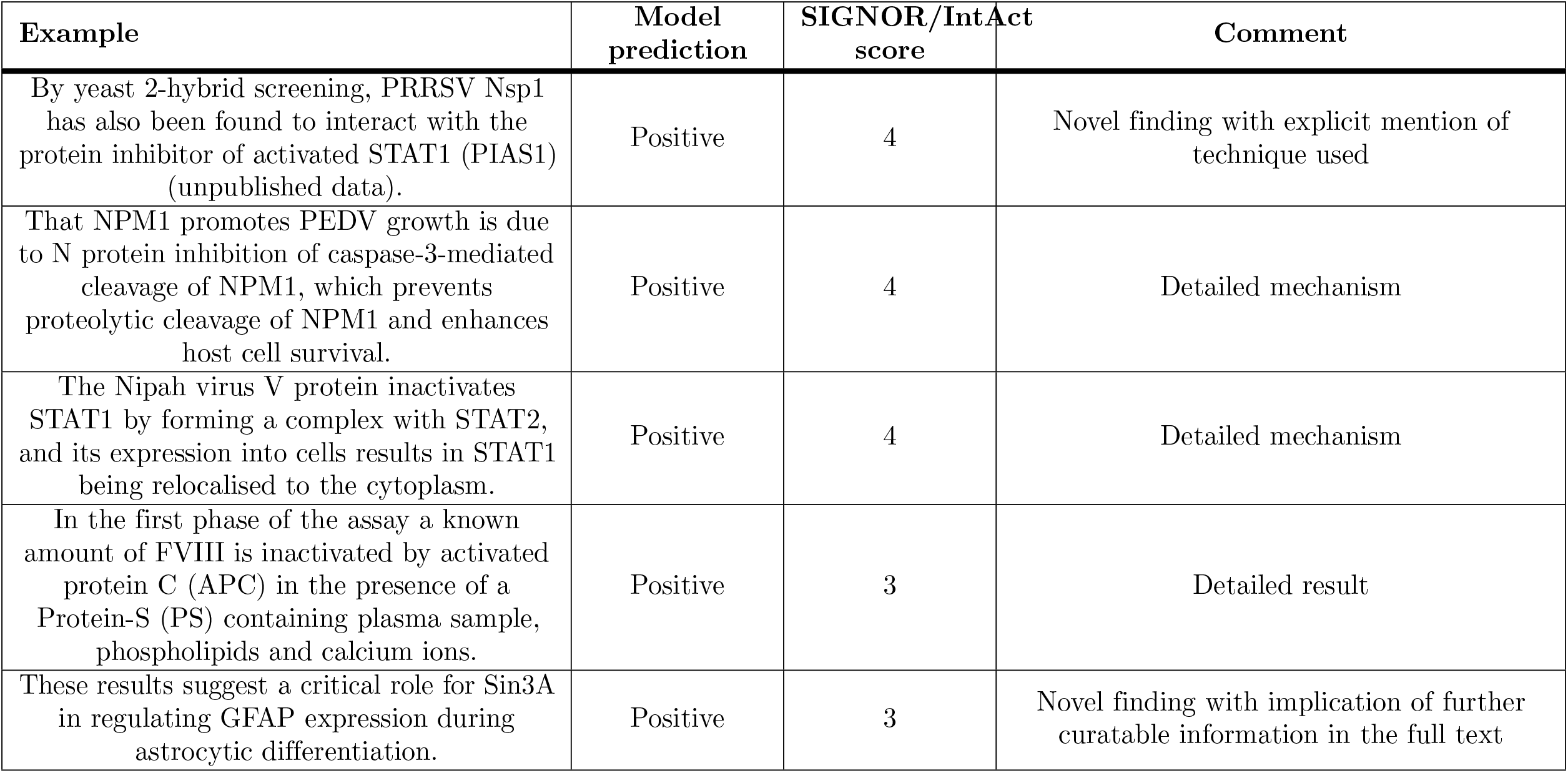
Illustrative high grade SIGNOR/IntAct sentences.

## Conclusion

Various factors affect the quality and cost efficiency of training data for relationship extraction from text in the life sciences. Our key findings are summarised below.

- Named entity recognition provides a useful starting point for curation, as well as for identifying sentences to pass to the trained model at inference. It also allows for the automatic replacement of specific entities with generic tokens, preventing the model from simply remembering protein pairs, which again applies to both training and inference.
- Inter-annotator agreement is essential for high quality data. Life science text is deeply complex and individuals regularly disagree when asked to classify content. In the case of PPIs, a voting mechanism with three independent curators dissolved most of these conflicts.
- Semisupervision may be a valuable preprocessing step where significant class imbalances are present in the data available for curation. Some entity relationships seem to be more amenable to this approach than others.
- Rules-based approaches offer interpretable results, but require careful manual tuning to balance bias and variance. While rules may be indefinitely tuned, in practice it is challenging to achieve comparable efficacy to deep learning results.
- Deep learning methods appear to be capable of some degree of abstraction when bootstrapped with data collected from rules-based methods. This seems to be particularly true where the rules in question account for grammatical dependency as opposed to pure named entity recognition.
- BioBERT is capable of achieving strong results with relatively few training samples. 500 sentences per class seems to be a reasonable target for an initial round of curation, at which point the accuracy curve should be assessed.
- Capturing examples that are unclear in a separate class may allow for training two models with emphasis either on recall or precision.
- If a model is intended for use in multiple different settings, it is possible to fine tune a recall-oriented model to more specific criteria with relatively few training samples.

The SIGNOR/IntAct curation validated the use of relationship extraction models to streamline the identification of protein pairs for curation. However, it also illustrated the importance of further fine tuning to target specific subcategories within the scope of all sentences containing PPIs, and provided ideas for future work. In particular we feel that the feasibility of developing a model for detecting novel findings warrants investigation. This could then be used in combination with the PPI model, or any other relationship extraction model, to filter results such that only sentences describing the initial identification of a relationship were targeted. These would be more likely to be accompanied by the evidence and quantitative metrics required for comprehensive database curation. Moderate gains were made by further fine tuning of the model using a small training set curated according to the different criteria of SIGNOR/IntAct, illustrating the potential for a model with broad criteria to be fine tuned to more specific use cases.

## Supporting information

**S1 Appendix. PPI curation guidelines**. The full set of high level guidelines provided to curators.

**S2 Appendix. Biomolecular interaction terms**. The full set of terms used in the rules-based methods to identify candidate sentences containing two genes/proteins and one or more molecular bioverbs.

**S3 Appendix. PPI technology terms**. The full set of terms used to filter documents for the initial SIGNOR/IntAct curation on CORD19 data.

**S4 Appendix. SIGNOR classifications**. The definitions for SIGNOR’s sentence level classification effort.

## Acknowledgments

We extend our thanks to Pablo Millan of the European Bioinformatics Institute for some initial discussions, curation efforts and for referring us to SIGNOR.

